# Clonal Expansion of Alveolar Fibroblast Progeny Drives Pulmonary Fibrosis

**DOI:** 10.1101/2025.05.31.657194

**Authors:** Christopher Molina, Tatsuya Tsukui, Imran Khan, Xin Ren, Wenli Qiu, Michael Matthay, Paul Wolters, Dean Sheppard

## Abstract

Pulmonary fibrosis has been called a fibroproliferative disease but the functional importance of proliferating fibroblasts to pulmonary fibrosis has not been systematically examined. In response to alveolar injury, resting alveolar fibroblasts differentiate into fibrotic fibroblasts that express high levels of collagens. However, what role, if any, proliferation plays in the accumulation of fibrotic fibroblasts remains unclear. Through EdU incorporation, genetic lineage tracing, and single cell RNA sequencing, we resolve the proliferation dynamics of lung fibroblasts during post-injury fibrogenesis. Our data show substantial DNA replication in progeny of alveolar fibroblasts in two models of pulmonary fibrosis. By genetically labeling individual cells, we observe clonal expansion of alveolar fibroblast descendants principally in regions of fibrotic remodeling. The transcriptome of proliferating fibroblasts closely resembles that of fibrotic fibroblasts, suggesting that fibroblasts can first differentiate into fibrotic fibroblasts and then proliferate. Genetic ablation of proliferating fibroblasts and selective inhibition of cytokinesis in alveolar fibroblast descendants significantly mitigates pulmonary fibrosis and rescues lung function. Furthermore, fibroblasts in precision-cut lung slices from human fibrotic lungs exhibit higher proliferation rates than those in non-diseased lungs. This work establishes fibroblast proliferation as a critical driver of pulmonary fibrosis and suggests that specifically targeting fibroblast proliferation could be a new therapeutic strategy for fibrotic diseases.

## INTRODUCTION

Idiopathic Pulmonary Fibrosis (IPF) has been characterized as a progressive fibroproliferative disease. However, the evidence supporting an important role for fibroblast proliferation in human fibrotic lungs is controversial. Numerous studies have reported both enhanced (1–9) and diminished (10–19) proliferative capacities of fibroblasts derived from human fibrotic lungs. These discrepancies likely stem from variations in culture conditions, cell purification techniques, and heterogeneity in the distribution of lesions typically encountered in fibrotic lungs (20, 21).

Given the limited effectiveness of current therapies for IPF, it is crucial to deepen our understanding of the mechanisms underlying the progressive nature of this disease. Progressive collagen deposition in IPF is thought to originate from differentiated Cthrc1^+^ fibrotic fibroblasts (22, 23). These Cthrc1^+^ fibroblasts can assemble into pathologic aggregates called fibroblastic foci (22, 24); and increasing number of these foci (25, 26) and higher bulk Cthrc1 expression correlate with more severe disease (27). These foci exhibit significant variability in size, which could indicate temporal growth, with the smallest foci occurring at the most recent sites of injury (28), then dynamically expanding outward forming an invasive fibrotic front (29). However, the contribution of fibroblast proliferation to the formation of these pathologic fibroblast aggregates remains unresolved, with studies variably reporting either enriched (30–32) or diminished (29, 33–37) staining of proliferation markers in fibroblastic foci.

It is not possible to directly examine the functional importance of fibroblast proliferation in human pulmonary fibrosis. Consequently, extensive research has been directed towards understanding fibroblast proliferation in rodent models. However, these rodent studies have also yielded conflicting results, with some studies suggesting that proliferating mesenchymal cells contribute to lung fibrosis (38–50), while other studies suggest a minimal role for mesenchymal cell proliferation (51–53). These inconsistencies could be in part explained by limited understanding of the heterogeneity of fibroblast populations in the lungs and the absence of reliable tools to label and manipulate specific subsets of fibroblasts.

To date, no studies have definitively demonstrated that in vivo fibroblast proliferation is a significant driver of pulmonary fibrosis. One recent study utilizing Pdgfrα-CreER in an influenza model observed a reduction in both lung injury severity and dysplastic epithelial regeneration when Ect2 was deleted to inhibit cytokinesis in Pdgfrα^+^ fibroblasts (54), but did not directly examine fibrosis as an endpoint. Ect2, a Rho GDP exchange factor (RhoGEF), plays an important role in both cytokinesis and cell migration (55) so that study did not formally determine whether the effect on dysplastic regeneration was due to inhibition of cytokinesis or fibroblast migration. Also, since Pdgfrα labels a heterogenous group of lung fibroblasts, including normal alveolar and adventitial fibroblasts, that study did not definitively determine which population was responsible for the effects observed.

Recent scRNA data has identified Scube2 as a marker of normal alveolar fibroblasts in mice (22) and demonstrated that Scube2^+^ alveolar fibroblasts directly support the alveolar type II stem cell niche (23). In response to alveolar insults that lead to the development of pulmonary fibrosis, Scube2^+^ alveolar fibroblasts are the primary cell of origin for multiple emergent fibroblast subsets including, fibrotic, inflammatory, stress activated and proliferating fibroblast subsets (23). Based on these results, we recently developed tools to precisely label resting alveolar fibroblasts (Scube2-CreER) and induced fibrotic fibroblasts (Cthrc1-CreER).

In this study, we use these tools, together with efficient (Rosa26-lox-stop-lox TdTomato) and inefficient (Brainbow2.1/+ confetti) lineage tracing, EdU labeling and genetic deletion of Ect2 (which inhibits cytokinesis and migration) and Esco2 (which impairs chromatin cohesion and thus leads to death of proliferating cells) to directly examine the extent, functional significance and molecular characteristics of fibroblast proliferation in two distinct models of pulmonary fibrosis induced by silica or bleomycin. We also reanalyze scRNA sequencing data from human lungs and analyze fibroblast proliferation in human precision cut lung slices to assess the relevance of murine findings to pulmonary fibrosis in humans. We find substantial fibroblast proliferation in both murine models and show that the predominant population of proliferating fibroblasts are Cthrc1^+^ fibrotic fibroblasts derived from Scube2^+^ lineage-traced alveolar fibroblasts. These descendants of alveolar fibroblasts clonally expand, forming pathologic fibroblast clusters in regions of fibrotic remodeling. We show that selective inhibition of cytokinesis or migration (by deletion of Ect2) or induction of cell death in proliferating cells (by deleting Esco2) in cells derived from alveolar fibroblasts diminishes the formation of pathologic fibroblast clusters, reduces collagen accumulation, and rescues organ function in two in vivo models of pulmonary fibrosis. Further, we show that human lung fibroblasts in fibrotic lung slices proliferate more extensively than fibroblasts in normal human lung slices and that in scRNA sequencing data from human lungs CTHRC1^+^ fibroblasts have the highest rate of expression of genes associated with cell cycle progression. These findings strongly suggest that, in response to fibrotic alveolar injury, alveolar fibroblasts can differentiate into fibrotic fibroblasts and then proliferate. This proliferation significantly contributes to the formation of fibroblast aggregates, the accumulation of excess collagen and the subsequent impairment of gas exchange and lung function. Since fibroblast proliferation also appears to occur in fibrotic fibroblasts in human fibrotic lungs, these results suggest that targeted inhibition of fibroblast proliferation could be a viable strategy for the treatment of pulmonary fibrosis.

## RESULTS

### Evaluating Lung Fibroblast Proliferation Dynamics in Two In Vivo Models of Pulmonary Fibrosis

Given the pivotal role of fibroblasts in lung scarring, we postulated that fibroblast proliferation is a fundamental mechanism in fibrotic lung remodeling. To this end, we analyzed the proliferation dynamics of lineage-negative (CD45^-^, CD31^-^, Epcam^-^, Mcam^-^) lung fibroblasts in two in vivo models of lung fibrosis. We employed adult C56BL/6 wild-type mice, challenging them with a single intra-airway dose of either bleomycin or silica, followed by daily injections of Ethinyl Deoxyuridine (EdU) for four days before harvesting at intervals from days 0 to 16 post-injury. This allowed us to quantify the rolling average of proliferating fibroblasts at these time points (Figure S1A-S1B). Our findings indicate significant rates of EdU incorporation in both models, with peak effect occurring between days 4-8 post-injury in silica-treated lungs and days 8-12 in bleomycin-treated lungs (Figure S1C). In both models a substantial fraction of fibroblasts had incorporated EdU during all time intervals studied.

### Origin of Proliferating Fibroblasts in Lung Fibrosis Models

Recent studies demonstrated a progressive increase in a population of Cthrc1^+^ fibrotic fibroblasts in response to bleomycin and showed that Scube2^+^ alveolar fibroblasts are the major progenitor source of these cells(23). However, whether aggregates of fibrotic fibroblasts simply emerge from existing alveolar fibroblasts or expand due to proliferation remains unclear. To begin to delineate the role of proliferation, we utilized Scube2-CreER; Rosa26-Ai14 mice to label alveolar fibroblasts with tdTomato, induced fibrosis using bleomycin or silica, and administered EdU in the drinking water over the first 21 days post-injury (Figure 1A). Remarkably, an average of 73.9 (± 8.0)% of EdU-labeled fibroblasts were tdTomato-labeled (Figure 1B), closely mirroring the 78.1% labeling efficiency of alveolar fibroblasts by Scube2-CreER (Figure 1C) (22), suggesting that vast majority of fibroblasts incorporating EdU in each fibrosis model were derived from alveolar fibroblasts.

**Figure 1:**
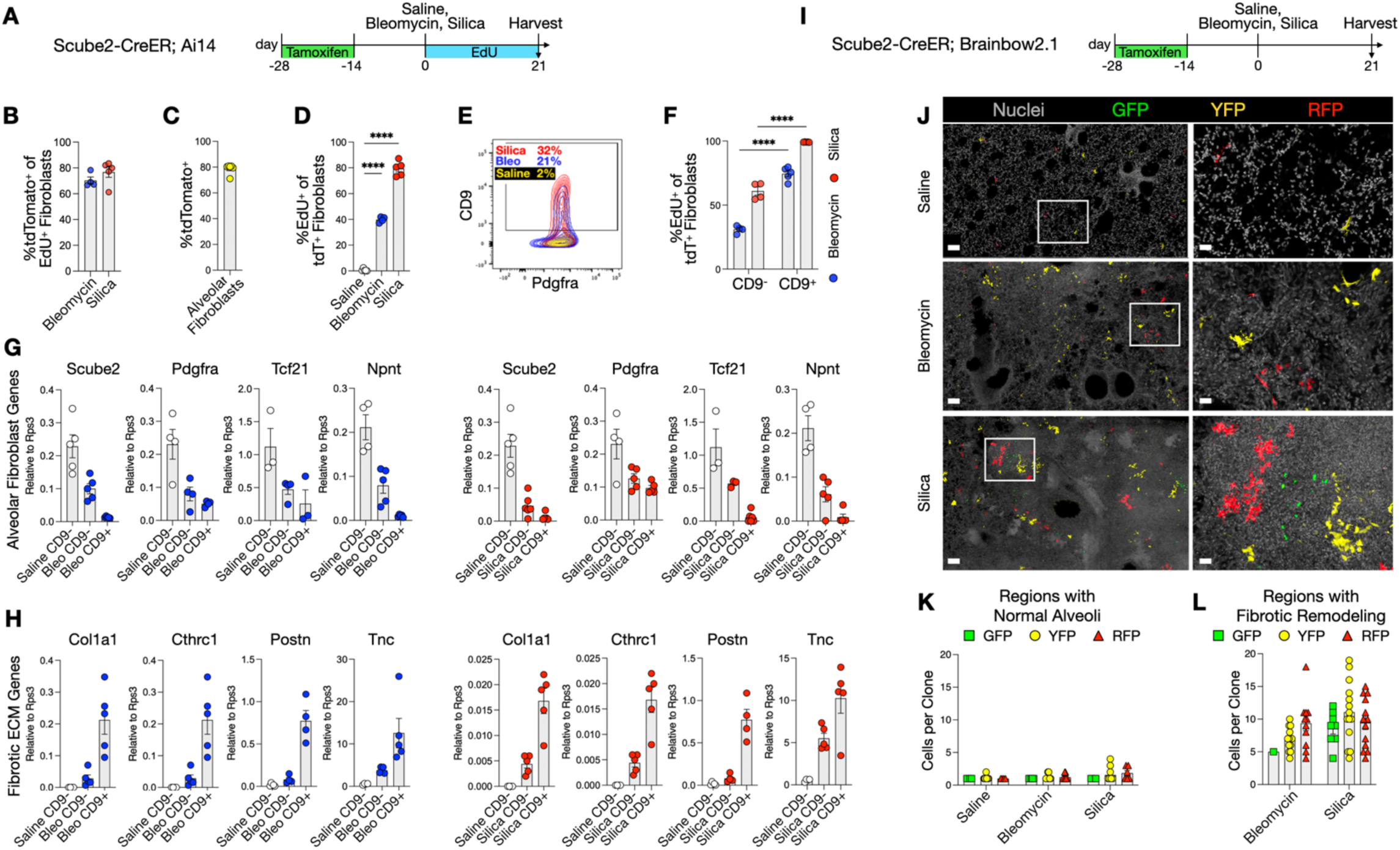
Alveolar Fibroblast Descendants Proliferate in Regions of Fibrotic Remodeling and Adopt a Profibrotic Phenotype. (**A**) Alveolar fibroblasts in Scube2-CreER/Rosa26-Ai14 mice were labeled with tdTomato via tamoxifen injection, subjected to bleomycin or silica challenge, then treated with EdU in the drinking water to label proliferating cells. On day 21, tdTomato^+^ fibroblasts were quantified and sorted via FACS for qPCR. (**B**) In bleomycin-treated lungs, 70.3 (± 5.4) % of all lineage-negative fibroblasts that proliferated at least once were tdTomato^+^, compared to 76.8 (± 9.0) % in silica-treated lungs. (**C**) In saline-control mice, the labeling efficiency of alveolar fibroblasts by Scube2-CreER was 78.1 (±3.9) %. (**D**) 39.9 (± 2.2) % of tdTomato^+^ fibroblasts proliferated at least once in bleomycin-treated lungs, compared to 79.4 (± 5.7) % in silica-treated lungs, and 1.3 (± 0.9) % in saline-control lungs. (**E**) CD9 upregulation was noted in 20-30% of tdTomato^+^ fibroblasts under fibrotic conditions. (**F**) CD9^+^ tdTomato^+^ fibroblasts showed higher proliferation rates compared to CD9^-^ tdTomato^+^ fibroblasts in bleomycin and silica-treated lungs. (**G-H**) qPCR on sorted cells revealed changes in gene with CD9^+^ fibroblasts, preferentially downregulating genes associated with quiescent alveolar fibroblasts (**G**) and upregulating fibrotic ECM genes (**H**). (**I**) Sparse labeling with fluorescent GFP, YFP, or RFP in Scube2-CreER; Rosa26-Brainbow2.1/+ mice allowed for tracking of fibroblast clonal expansion. (**J-L**) Confocal microscopy of cleared lung sections from treated mice revealed cellular expansions in fibrotic areas, quantified by cellular density per clone in fibrotic versus normal regions. Statistical analysis for (D) and (F) was performed with 2-way ANOVA; error bars represent SEM, with significant proliferative differences denoted (****p<0.0001).

We observed that 39.9% of tdTomato-labeled fibroblasts in bleomycin-injured lungs and 79.4% in silica-injured lungs were EdU labelled (Figure 1D). We previously showed that the cell surface marker, CD9, is not normally expressed by alveolar fibroblasts, but at 21 days after bleomycin treatment CD9 expression is specifically induced in fibrotic fibroblasts (22). We therefore sorted tdTomato positive cells for CD9 expression at the 21-day endpoint of EdU labeling and compared EdU incorporation if CD9^+^ cells compared to CD9^-^ cells. In both models, a higher fraction of CD9^+^ cells had undergone at least one round of DNA replication (as detected by EdU labeling) with values of 74.5% after bleomycin and 99.2% after silica (Figures 1E, 1F). Quantitative PCR of sorted cells revealed that CD9^+^ tdTomato^+^ fibroblasts significantly downregulated alveolar fibroblast genes (Scube2, Pdgfra, Tcf21, Npnt) and upregulated fibrotic extracellular matrix genes and markers of fibrotic fibroblasts (Col1a1, Cthrc1, Postn, Tnc), indicating their transition from alveolar to fibrotic fibroblasts (Figure 1G, 1H). These results suggest that proliferating fibroblasts predominantly arise from alveolar fibroblast progenitors, and that most CD9^+^ fibrotic fibroblasts emerged from cells that had undergone at least one round of DNA replication.

### Spatial Distribution of Fibroblast Proliferation in Fibrotic Lungs

To further elucidate the spatial distribution and extent of fibroblast proliferation in fibrotic lungs, we utilized Scube2-CreER; Brainbow2.1/+ confetti mice (74). After administering tamoxifen to induce sparse labeling of alveolar fibroblasts and their descendants with GFP, YFP, or RFP, we subjected the mice to challenges with bleomycin, silica, or saline (Figure 1I). Confocal imaging of cleared thick sections demonstrated clearcut clonal expansion of alveolar fibroblast progeny and this expansion predominantly occurred in areas of dense fibrotic remodeling (Figures 1J-L). These results support the conclusion that EdU incorporation in alveolar fibroblast progeny is a true marker of cell proliferation and identify regions of dense fibrotic remodeling as the principal site of clonal expansion in both fibrosis models.

### Heterogeneity of Proliferating Fibroblasts in Lung Fibrosis

Generally, it has been thought that fibroblasts proliferate before they differentiate into ECM-producing fibroblasts (56–59). To explore the molecular phenotype of proliferating fibroblasts, we leveraged our previously published scRNA-seq dataset (23), which highlighted the emergence of distinct fibroblast subpopulations in bleomycin-induced fibrotic lungs — fibrotic, inflammatory, and proliferative fibroblasts (Figure 2B). Our re-analysis of the proliferative fibroblast cluster (Figure 2C, Supplemental Figure 3A), confirmed that most of the proliferating fibroblasts originated from Scube2-CreER alveolar fibroblasts, aligning with our continuous EdU labeling results (Figure 1B). Notably, the largest fraction of proliferating fibroblasts were present at day 7 post-injury (Figure 2D), which we verified using 24-hour EdU pulse injections in Scube2-CreER; Ai14 mice (Supplemental Figure 2).

**Figure 2:**
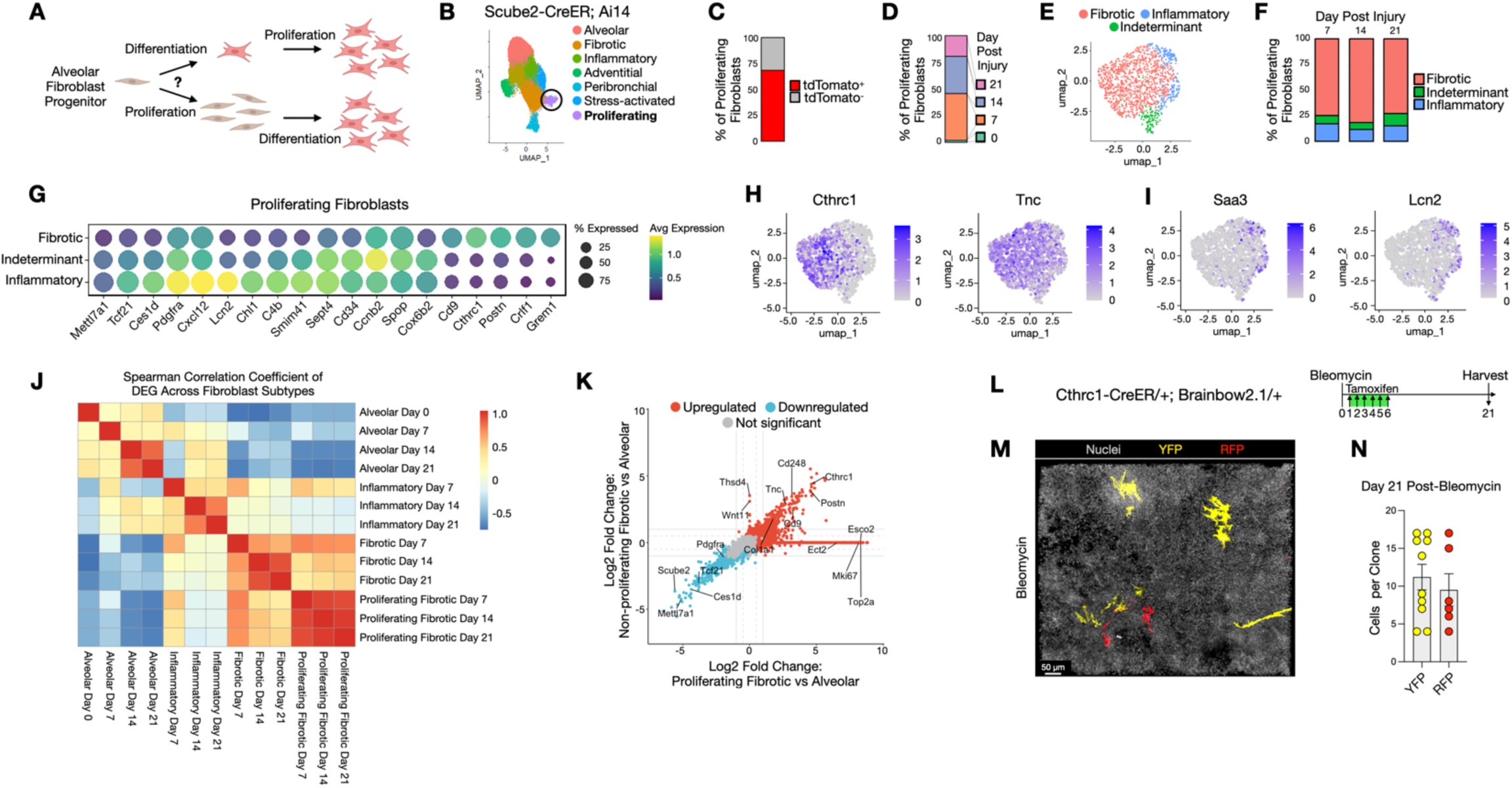
Cthrc1+ Fibrotic Fibroblasts Are the Dominant Proliferating Fibroblast Subtype in Fibrotic Lungs. (**A**) Potential pathways of fibroblast proliferation and differentiation in fibrotic lungs (**B**) UMAP visualization from Tsukui et al., 2024, showing fibroblasts harvested from Scube2-CreER; R26-Ai14 mice at days 0, 7, 14, and 21 post-bleomycin injury. The proliferating fibroblast cluster was subsetted and reclustered for downstream analysis. (**C**) Most proliferating fibroblasts were lineage-labeled by tdTomato. (**D**) Fibroblast proliferation was maximally observed at day 7 post-bleomycin injury. (**E**) Reclustering identified three distinct proliferating fibroblast subpopulations: fibrotic, inflammatory, and indeterminate. (**F**) Fibrotic fibroblasts consistently emerged as the predominant proliferating type across all time points. (**G**) Bubble plot and feature plots (**H-I**) displaying marker gene expression profiles across subtypes. (**J-K**) Spearman correlation and scatter plot analyses confirmed the transcriptomic similarities between proliferating and non-proliferating fibrotic fibroblasts. Dashed grey line ±0.5 average log2fc; solid grey line ±1 average log2fc. (**L-N**) Longitudinal tracking of Cthrc1+ fibrotic fibroblast clonal expansion in bleomycin-treated lungs, visualized using cleared thick sections by confocal microscopy, demonstrating significant cellular proliferation within fibrotic regions.

Further re-clustering of the proliferating fibroblast population revealed three transcriptionally distinct subtypes: fibrotic, inflammatory, and a smaller indeterminate group (Figures 2E-I). The inflammatory subtype was characterized by the expression of inflammatory genes such as *Saa3, Lcn2, Cxcl12*, and *C4b* (Figures 2G, 2I), which overlap with markers of the “transition” fibroblasts described by Mayr/Sengupta et al. and Konkimalla et al. (Supplemental Figure 3) (60, 61). The proliferating fibrotic subtype predominantly expressed genes that are typically upregulated in fibrotic fibroblasts, such as *Cthrc1, Tnc, Postn*, and *Grem1* (Figures 2G, 2H) (23, 24, 27). Surprisingly, the Cthrc1^+^ fibrotic fibroblast was the dominant proliferative subpopulation across all time points analyzed (Figure 2F).

To delineate how proliferative fibrotic fibroblasts compare with non-proliferative fibrotic fibroblasts at the global transcriptome level, we performed a Spearman correlation analysis across fibroblast subtypes, which confirmed that proliferative fibrotic fibroblasts closely resemble non-proliferative fibrotic fibroblasts (Figure 2J). A comparison of the differential gene expression profiles between these two states showed nearly identical patterns in their downregulation of alveolar fibroblast genes such as *Scube2, Pdgfra, Tcf21, Mettl7a1*, and upregulation of CD9 and fibrotic ECM genes such as *Col1a1, Cthrc1, Postn*, and *Tnc* (Figure 2K). Unique markers distinguishing the proliferative from non-proliferative fibrotic fibroblasts predominantly included cell cycle genes like *Mki67, Top2a, Esco2*, and *Ect2* (Figure 2K).

To further test the ability of differentiated fibrotic fibroblasts to proliferate, we treated Cthrc1-CreER; Brainbow2.1/+ mice with bleomycin then administered tamoxifen on days 1-6 to sparsely label emerging fibrotic fibroblasts with either RFP or YFP (Figure 2L). By day 21 post-injury, clonal expansion of Cthrc1-CreER-labeled fibroblasts was evident (Figure 2N), mirroring patterns observed in bleomycin-treated Scube2-CreER; Brainbow2.1/+ mice (Figure 1L), confirming that alveolar progeny that have already differentiated into fibrotic fibroblasts can undergo substantial clonal expansion (Figure 2A).

### Functional Role of Proliferating Fibroblasts in Fibrotic Lung Injury Models

To assess the effects of eliminating proliferating fibroblasts, we treated Scube2-CreER; Ai14; Esco2-fl/fl mice with tamoxifen, which activated tdTomato labeling and deleted the Esco2 gene in alveolar fibroblasts and their descendants. Esco2 is a crucial element of the chromatin cohesion complex, and loss of Esco2 in proliferating cells results in cell apoptosis (62). Thus, deleting Esco2 allows targeting of proliferating fibroblasts for apoptosis during lung fibrogenesis induced by bleomycin and silica (Figure 3A). At day 28, whole lung imaging revealed marked reductions in dense tdTomato^+^ fibroblast aggregates in Esco2-deficient mice compared to controls in both bleomycin and silica models (Figure 3B). Flow cytometry quantification confirmed a decrease in total tdTomato^+^ fibroblasts in the treated Scube2-CreER; Ai14; Esco2-fl/fl mice (Figure 3C). Assessments of total lung hydroxyproline content indicated a substantial decrease in fibrosis severity in Esco2-fl/fl mice (Figures 3D, 3E), while pulse oximetry measurements demonstrated improved lung oxygenation in Esco2-fl/fl mice (Figures 3F, 3G).

**Figure 3:**
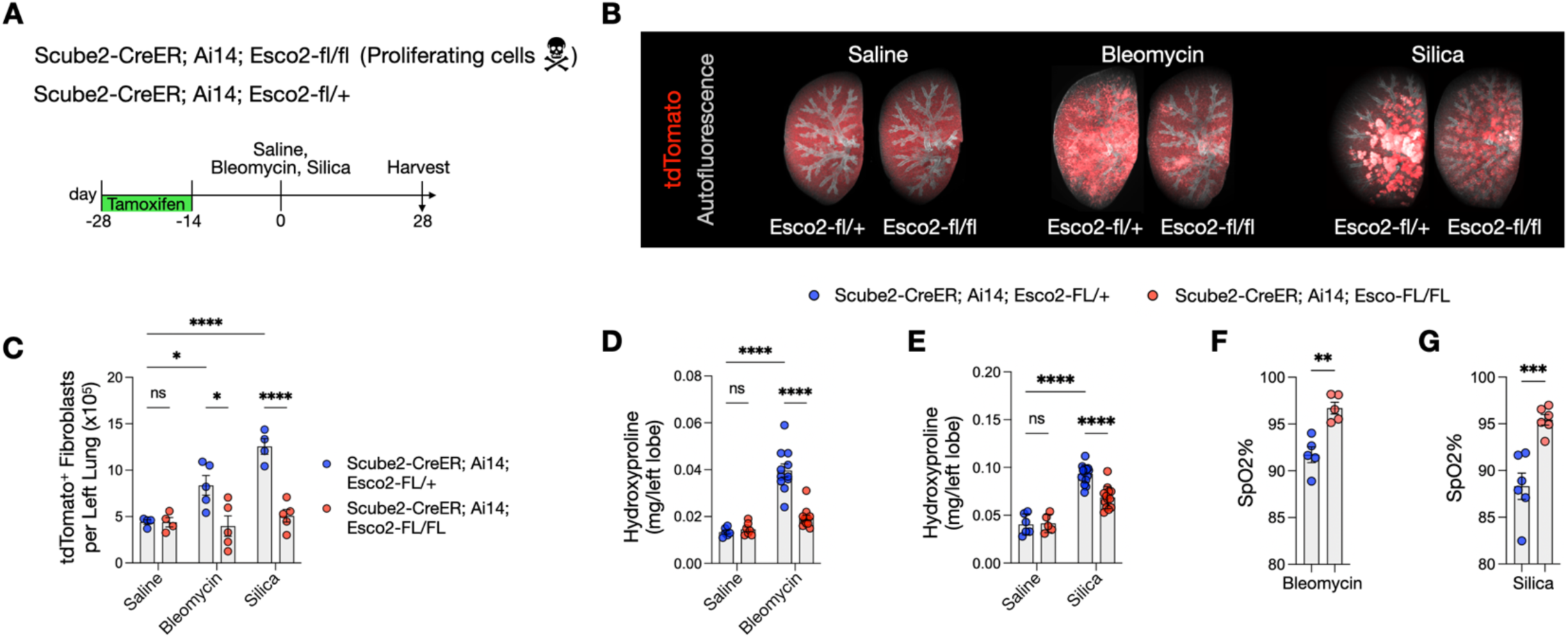
Genetic Deletion of Proliferating Fibroblasts Reduces Pulmonary Fibrosis and Restores Lung Function. (**A**) Tamoxifen was administered to label alveolar fibroblasts with tdTomato and delete Esco2 in Scube2-CreER; Ai14; Esco2-fl/fl mice, targeting proliferating progeny for ablation. These mice and Scube2-CreER; Ai14; Esco2-fl/+ controls underwent bleomycin, silica, or saline challenges, with analysis at day 28. (**B**) Maximum intensity projections of cleared right upper lobes, imaged by lightsheet microscopy, highlighting tdTomato^+^ cells in red against grey-pseudocolored autofluorescent airways. (**C**) Flow cytometric analysis revealed a reduction in the total number of tdTomato^+^ fibroblasts in fibrotic lungs of Esco2-fl/fl mice; n=8 saline, n=11 bleomycin, n=9 silica. (**D, E**) Hydroxyproline assays demonstrated reduced lung fibrosis in both bleomycin and silica-treated Esco2-fl/fl lungs; n=22 saline, n=23 bleomycin, n=29 silica. (**F, G**) Improved lung oxygenation was quantified via pulse oximetry in bleomycin and silica-treated Esco2-fl/fl mice; n=10 bleomycin, n=12 silica. Statistical analyses for (C), (D), and (E) were performed using 2-way ANOVA with Tukey’s multiple comparisons test; (F) and (G) were analyzed by unpaired parametric t-tests. Error bars represent SEM. * p<0.05, ** p<0.01, *** p<0.001, **** p<0.0001.

To explore the consequences of selectively inhibiting fibroblast proliferation without killing, we deleted Ect2 in Scube2-CreER; Ai14; Ect2-fl/fl mice, to impair cytokinesis (55) in tdTomato^+^ alveolar fibroblasts under similar conditions (Figure 4A). At day 28 post-injury, whole lung imaging showed reductions in tdTomato^+^ fibroblast aggregates in the Ect2-deficient group (Figure 4B). This was corroborated by flow cytometry data showing a reduction in the total number of tdTomato^+^ fibroblasts (Figure 4C), and hydroxyproline assays confirming decreased fibrosis in these mice (Figure 4D). Pulse oximetry revealed significant improvements in oxygenation in the Ect2 model, emphasizing the beneficial effects of inhibiting fibroblast proliferation on overall lung health (Figures 4E, 4F). Together, these loss of function experiments demonstrate that fibroblast proliferation worsens fibrosis severity and lung function in two in vivo models of pulmonary fibrosis.

**Figure 4:**
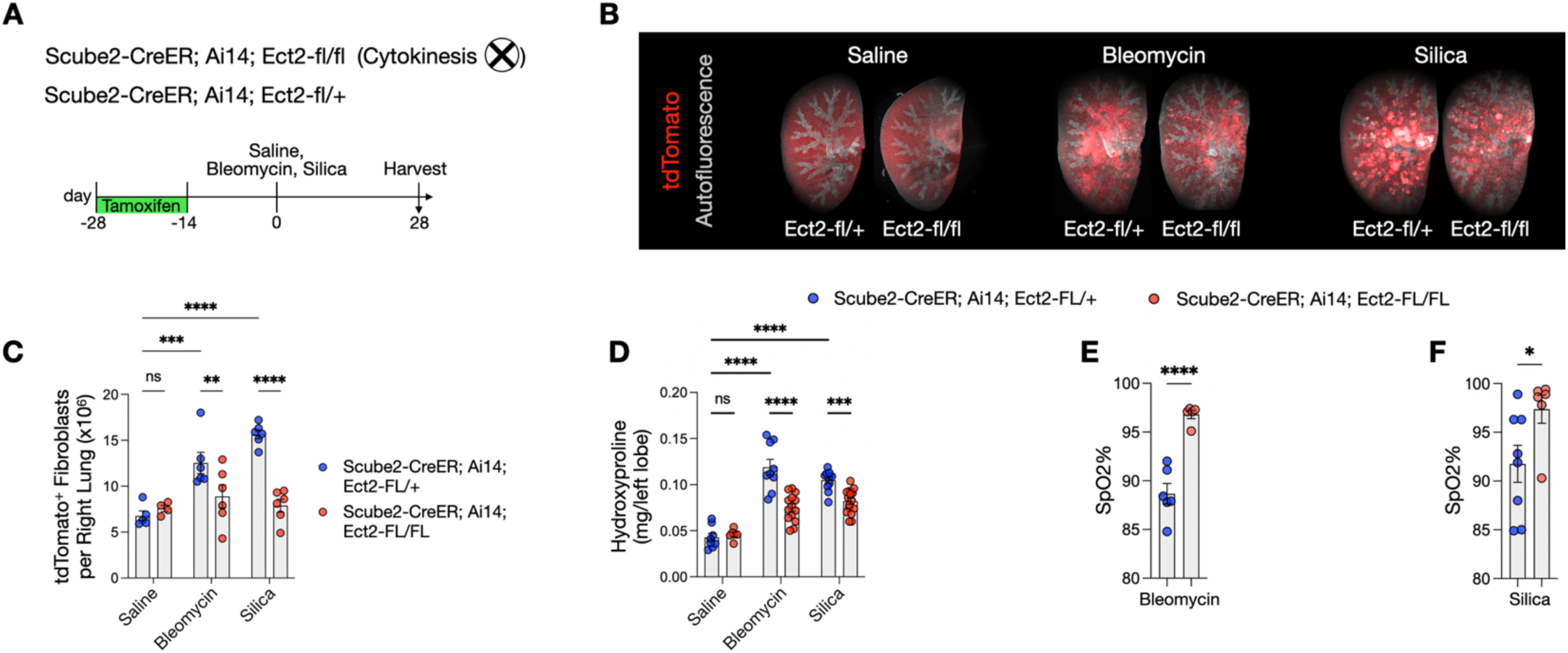
Genetic Inhibition of Fibroblast Proliferation Ameliorates Lung Fibrosis and Restores Lung Function. (**A**) Treatment of Scube2-CreER; Ai14; Ect2-fl/fl mice with tamoxifen enabled both tdTomato labeling of alveolar fibroblasts and deletion of Ect2, allowing targeted inhibition of cytokinesis in proliferating fibroblasts. These and Scube2-CreER; Ai14; Ect2-fl/+ control mice were subjected to challenges with bleomycin, silica, or saline, and analyzed at day 28. (B) Maximum intensity projections of cleared right upper lobe sections visualized via lightsheet microscopy show tdTomato^+^ cells in red and autofluorescent airways pseudocolored in grey. (C) Quantification of tdTomato^+^ fibroblasts by flow cytometry demonstrated a reduction in proliferative fibroblasts in Ect2-fl/fl mice; n=9 saline, n=12 bleomycin, n=12 silica. (D) Hydroxyproline assays indicate decreased fibrosis in Ect2-fl/fl mice challenged with bleomycin and silica; n=14 saline, n=23 bleomycin, n=26 silica. (E, F) Pulse oximetry measurements showed enhanced lung oxygenation in Ect2-fl/fl mice treated with bleomycin and silica; n=10 bleomycin, n=14 silica. Analyses for (C) and (D) utilized 2-way ANOVA with Tukey’s correction for multiple comparisons; (E) and (F) were assessed by unpaired parametric t-tests. Error bars denote SEM. * p<0.05, ** p<0.01, *** p<0.001, **** p<0.0001.

### Diversity and Dynamics of Proliferating Fibroblasts in Human Fibrotic Lungs

We hypothesized that the proliferative responses of fibroblasts in murine fibrotic lungs may exhibit some similarities to those in human fibrotic lungs. Thus, we conducted EdU labeling and FACS quantification of proliferation rates in lineage-negative (CD45^-^, CD31^-^, EPCAM^-^, MCAM^-^) human lung fibroblasts derived from precision-cut lung slices (PCLS) of both non-diseased donors and fibrotic lung explants (Figure 5A). These data show that human lung fibroblasts in fibrotic lung PCLS similarly manifest enhanced proliferative capacity (Figure 5B).

**Figure 5:**
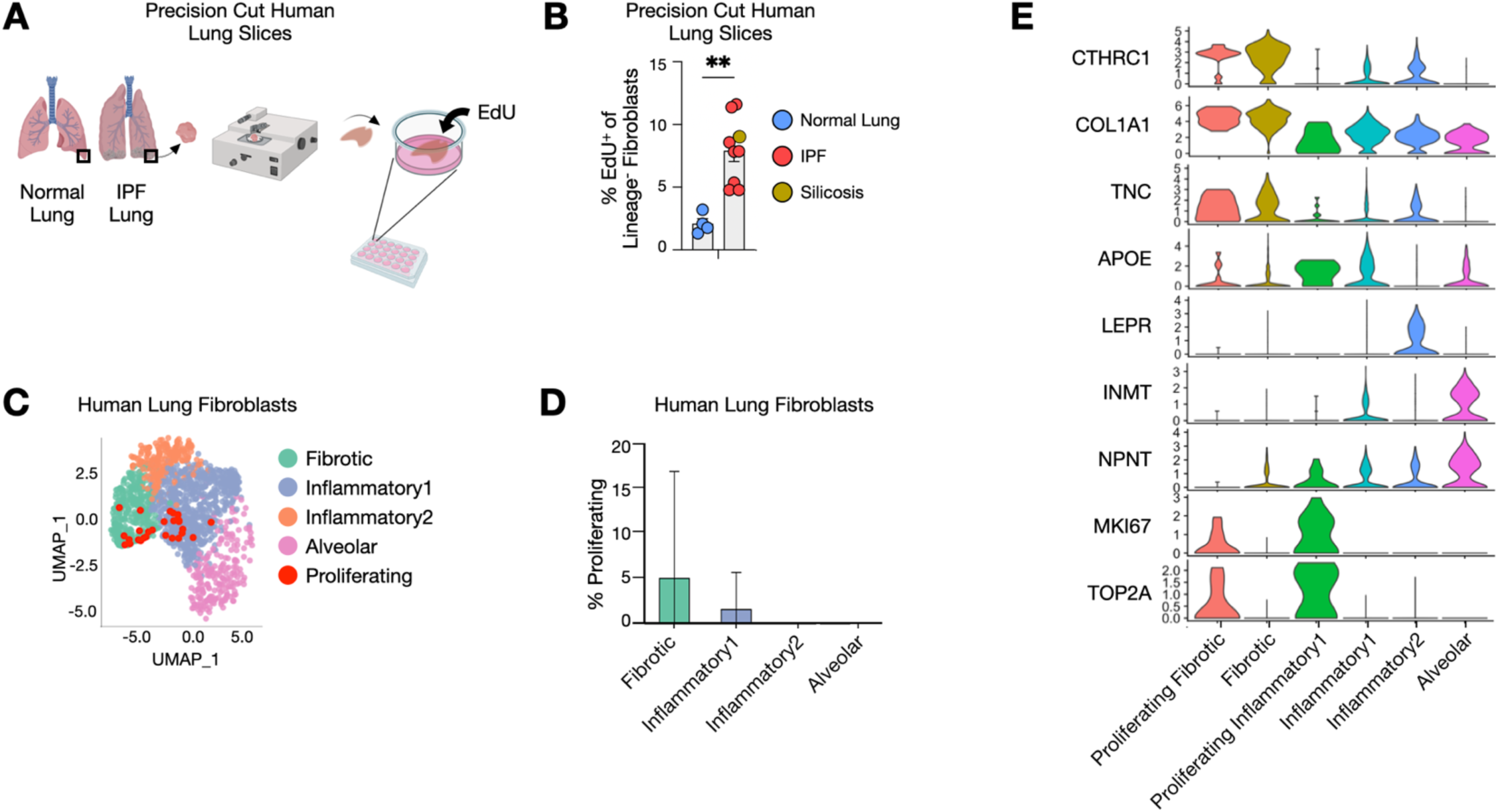
Heterogeneity of Proliferating Fibroblasts in Human Fibrotic Lungs. (**A-B**) Precision-cut human lung slices from both fibrotic and non-diseased donors were cultured in 1% FBS DMEM supplemented with EdU for two days then analyzed by flow cytometry; fibrotic lung donors, n=9; non-diseased lung donors, n=4. Statistical analysis was performed using an unpaired parametric t-test. ** p<0.01. Error bars represent SEM. (**C**) Re-analysis of the Habermann et al. IPF Cell Atlas by Tsukui et al. illustrating the presence of alveolar fibroblasts and three distinct pathologic fibroblast subtypes in human fibrotic lungs: fibrotic, inflammatory1, and inflammatory2. Cell cycle scoring revealed proliferating fibroblasts predominantly within the fibrotic and inflammatory1 fibroblast subpopulations. (**D**) Proportion of proliferating cells within each fibroblast subtype. (**E**) Violin plots delineating expression profiles of marker genes across fibroblast subtypes.

Building on our findings in murine models, we hypothesized that these proliferating fibroblasts in human fibrotic lungs could similarly adopt a profibrotic phenotype. To test this hypothesis, we first analyzed the transcriptomic diversity of proliferating fibroblasts using the human idiopathic pulmonary fibrosis (IPF) single-cell atlas by Habermann et al., reanalyzed by Tsukui et al., which revealed the emergence of three distinct pathologic fibroblast subpopulations in fibrotic lungs (Figure 5C) (23, 63). To identify proliferating fibroblasts, we employed cell cycle scoring within Seurat, utilizing DBSCAN clustering to establish optimal cutoffs for cells expressing high levels of G2M and S phase cell cycle genes (Supplemental Figure 4). Fibroblasts exhibiting elevated G2M or S scores were re-annotated as "Proliferating" and subsequently overlayed on UMAP plots of all alveolar and fibrosis-associated fibroblasts. This visualization demonstrated that all proliferating cells were contained with the fibrotic or inflammatory1 subtypes (Figure 5C). Quantitative analysis confirmed that fibrotic fibroblasts exhibited a trend of higher proliferation rates compared to their inflammatory1 counterparts (Figure 5D), consistent with observations in our bleomycin-induced murine model (Figure 2F). The expression profiles of key pathologic extracellular matrix genes, such as COL1A1, CTHRC1, and TNC, were notably elevated in both proliferating and non-proliferating fibrotic fibroblasts (Figure 5E), paralleling the gene expression trends noted in murine models (Figure 2K).

## DISCUSSION

In the current study, we used two different murine models of pulmonary fibrosis to demonstrate ongoing DNA replication (measured by EdU labeling) in two different murine models of pulmonary fibrosis. Using mice we have recently described that uniquely target normal alveolar fibroblasts, we show that a large fraction of the progeny of alveolar fibroblasts in both models have undergone at least one round of DNA replication during the first 21 days after initiation of the models and that DNA replication is enriched in progeny of alveolar fibroblasts that have differentiated into fibrotic fibroblasts, identified by induction of the cell surface protein CD9 and confirmed by qPCR of sorted cells. We further demonstrate that this increase in DNA replication is associated with true cell proliferation, measured by an increase in the total number of alveolar fibroblast progeny and with substantial clonal expansion, measured by counting colony numbers after inefficient labelling in Brainbow mice. Furthermore, we demonstrate that clonal expansion was largely restricted to regions of dense fibrotic remodeling.

By re-analysis of our own publicly available single cell RNA sequencing we found that the overwhelming majority of fibroblasts we previously characterized as actively proliferating demonstrate patterns of gene expression that otherwise overlap with the molecular phenotype of fibrotic fibroblasts, suggesting that the induction of proliferation may occur simultaneously with or following the induction of the fibrotic molecular phenotype. This interpretation is further supported by our observation that fibrotic fibroblasts labeled early after treatment with bleomycin using the Cthrc1-CreER line underwent a similar degree of clonal expansion to that we observed labeling normal alveolar fibroblasts.

Finally, we report that deleting Esco2 to kill proliferating alveolar fibroblast progeny or deleting Ect2 to inhibit cytokinesis in the same cells both caused a reduction in fibroblast expansion and led to protection from pulmonary fibrosis and the resultant hypoxemia caused by either bleomycin or silica, suggesting that fibroblast proliferation is functionally important in driving fibrosis. Our findings that proliferating fibroblasts are present even in end stage fibrotic human lungs, that these cells often have a fibrotic molecular phenotype, and that fibroblasts in PCLS from fibrotic human lungs are more proliferative than fibroblasts in PCLS from normal human lungs suggest that these findings could have clinical relevance.

Prior studies investigating the extent of fibroblast proliferation in pulmonary fibrosis models have yielded conflicting results, with some studies suggesting important contributions of mesenchymal cell proliferation to fibrosis development (38–50), whereas others cast doubt on this conclusion (51–53). However, a major consideration in studies of mesenchymal cell proliferation involves the challenge of cellular heterogeneity within the fibroblast population (64), with uncertainty as to whether increased cell numbers reflect a generalized proliferative response or a selective expansion of a subpopulation (65).

Mayr/Sengupta et al. reported proliferating transitional fibroblasts and proliferating myofibroblasts similar to the Saa3^+^ inflammatory fibroblasts and Cthrc1^+^ fibrotic fibroblasts we describe here (Supplemental Figure 3) (60). Valenzi et al. showed proliferating Cthrc1^+^ fibrotic fibroblasts in human systemic sclerosis fibrotic lungs similar to our findings in IPF (66). While it has generally been thought that fibroblasts proliferate before they differentiate into ECM-producing fibroblasts (20, 56–59), our lineage-tracing data show that lung fibroblasts can differentiate into Cthrc1^+^ fibrotic fibroblasts and then proliferate and raise the possibility that this trajectory could even be the most common course. Moreover, the discovery of proliferating Cthrc1^+^ fibrotic fibroblasts in other datasets of fibrosis in the skin, colon, liver, and heart (66–68) suggests that the therapeutic potential of selectively targeting proliferating fibroblasts may be broadly applicable to other fibrotic diseases.

We also show, for the first time, that proliferating fibroblasts are functionally important in driving lung fibrosis and impairing gas exchange. We used two orthogonal strategies, specifically killing proliferating progeny of alveolar fibroblasts by deleting Esco2 and inhibiting proliferation without inducing cell death by interfering with cytokinesis by deleting Ect2. Each of these strategies has its own weaknesses. Deletion of Esco2 would induce fibroblast apoptosis, which could itself induce secondary responses in nearby cells, thereby confounding the conclusion that the protective effects of this intervention were simply due to loss of proliferating fibroblasts. Ect2 is a Rho guanidine exchange factor that inhibits cytokinesis by blocking the critical role of Rho activation in this process. However, Rho is also an important regulator of cell migration, so it is not possible to explicitly exclude inhibition of fibroblast migration as an explanation for the functional benefit of Ect2 deletion. However, our findings that both strategies led to similar protection from pulmonary fibrosis and hypoxemia in two different fibrotic models provides strong support for the conclusion that proliferation of alveolar fibroblast progeny plays an important role in pulmonary fibrosis and resultant functional impairment. These data are supported by prior studies showing strategies that impair fibroblast proliferation in the fibrotic heart (69, 70) and liver (71) improve organ function.

Several important questions remain unanswered by the work described here. These include identification of the molecular drivers of fibroblast proliferation, whether these drivers are shared among models of pulmonary fibrosis and whether similar pathways drive fibrosis in mice and humans. The similarities and differences between the signals that drive proliferation and those that drive the fibrotic phenotype also remain to be elucidated. All these questions should be the focus of future research.

It is obviously challenging to extrapolate our findings from these two murine models to conclusions about the functional significance of fibroblast proliferation in human pulmonary fibrosis. We were encouraged to see that we could identify proliferating fibroblasts even in explants from patients with end stage lung disease and that these proliferating fibroblasts were enriched among fibrotic fibroblasts, similar to what we saw in our murine models. Our observation that fibroblasts in PCLS from fibrotic human lungs were more likely to proliferate than fibroblasts from PCLS of normal human lungs provided further support for the notion that fibroblast proliferation could be important in human pulmonary fibrosis. Testing this hypothesis directly will require the development of therapeutic interventions that specifically target proliferating fibroblasts in vivo, ideally without targeting proliferation in other cell types. Given the intractable nature and high mortality rate of pulmonary fibrosis, we think our results should at least encourage efforts to develop such strategies.

## METHODS

### Sex as a biological variable

Our study examined male and female animals, and similar findings are reported for both sexes.

### Mice and bleomycin/silica treatment

The following mouse strains were used for experiments: C56BL/6 wild type mice (Jax Strain #000664), Scube2-CreER mice (Sheppard Lab) (23), Cthrc1-CreER mice (Sheppard Lab) (23), R26-Ai14 (Jax #007914), Brainbow2.1/Confetti (Jax #017492), Esco2-flox (Jax #030199), Ect2-flox (gift from Alan Fields) (55). Sex-matched littermates aged 12-16 weeks old were used for experiments. Because male mice develop more severe fibrosis after bleomycin treatment (72), male mice were treated with 2.5 units/kg of bleomycin and female mice were treated with 3 units/kg of bleomycin in 75 µL of saline by oropharyngeal aspiration. For silica treatments, average cohort weights were used to deliver 400 mg/kg of silica to male mice and 450 mg/kg of silica to female mice in 75 µL of saline by oropharyngeal aspiration. Silica (MIN-U_SIL5, US Silica) was baked in hydrochloric acid at 110 degrees Celsius overnight, then washed in sterile saline as previously described (23). Scube2-CreER mice were injected intraperitoneally with 2 mg of Tamoxifen (Millipore Sigma #T5648) dissolved in olive oil (Millipore Sigma #O1514, 20mg/mL) once daily for 2 weeks followed by a two-week washout period prior to treatment with bleomycin or silica. Cthrc1-CreER mice were injected intraperitoneally with 2 mg of Tamoxifen dissolved in olive oil once daily for 6 days after bleomycin treatment.

### 5 ethynyl-2-deoxyuridine (EdU) Treatments

For continuous labeling experiments, EdU (ThermoFisher #A10044) was dissolved in distilled water (1 mg/mL) and changed every other day for 21 days. For all pulse labeling experiments, EdU was dissolved in sterile PBS (10mg/mL) using a 50-degree water bath, then diluted in sterile PBS to deliver 50 mg of EdU per kg of mouse body weight in 200 µL of PBS. To measure the rolling average of EdU uptake across a four-day period (Supplemental Figure 1), mice were injected intraperitoneally with EdU once daily for four days prior to harvest. To measure EdU uptake during the 24 hours prior to harvest, EdU was injected intraperitoneally once at 23 hours prior to harvest and again at 1 hour prior to harvest. For precision-cut lung slices, EdU was dissolved in DMEM/F12+GlutaMAX (ThermoFisher #10565018) and added to culture media at 20 µM concentration. Lung slices without EdU were used as negative controls for flow cytometry gating. For all experiments, EdU uptake was measured by flow cytometry using the Click-iT Plus EdU Alexa Fluor 647 Kit (ThermoFisher #C10634).

### Tissue Dissociation

Mouse lungs were dissociated as previously described (23). In brief, mouse lungs were perfused with PBS through the right ventricle then the left lung was minced with scissors. The tissue was suspended in 1 mL of protease solution [0.25% Collagenase A (Millipore Sigma), 1 U/mL Dispase (Millipore Sigma), 2,000 U/mL DNase I (Millipore Sigma) in Hanks’ Balanced Buffer Solution (ThermoFisher)], then transferred into a 24 well plate and placed in a 37-degree incubator for 60 minutes with trituration by micropipette every 20 minutes. The single cell suspension was filtered through a 70 µM cell strainer (BD Biosciences) washed with PBS, then resuspended in 1 mL of ACK red blood cell lysis buffer (Gibco) for 90 seconds, then washed with 10% FBS, and resuspended in 1% Bovine Serum Albumin (Fisher BioReagents). For dissociation of human lung tissue, 3 precision cut lung slices, (approximately 1 cm height x 1 cm width x 500 µm thick each) were minced with scissors then suspended in 3 mL of protease solution, with all other downstream steps being identical to that of the mouse lung dissociation.

### Flow Cytometry

Following lung tissue dissociation, 3 x 10^6^ cells were used for flow cytometry. Cells were incubated with primary-conjugated antibodies in 1% Bovine Serum Albumin PBS for 30 minutes on ice. CountBright Plus Ready Tubes (ThermoFisher #C40000) were used for absolute cell counting. The following anti-mouse reagents were used at 1:100 concentrations unless otherwise specified: Live Dead Stain [DAPI 0.1 µg/mL; Live-or-Dye 330/410 (1:500), Biotium), CD45 (clone 30-F11, BV786, BD Biosciences), CD31 (Clone 390, BV605, BD Biosciences), EPCAM (clone G8.8, PE, BV421 Biolegend), MCAM (clone ME-9F1, Alexa 488, Biolegend, CD9 (clone MZ3, APC/Fire 750, Biolegend). The following anti-human reagents were used at 1:100 concentrations unless otherwise specified Live Dead Stain (Live-or-Dye 330/410 (1:200), Biotium), CD45 (clone HI30, APC-Fire 750, Biotin, Biolegend), CD31 (clone WM59, BV605, BV786, Biotin, BD Biosciences), EPCAM (clone 9C4, PE, Biolegend), MCAM (clone P1H12, PE-Cy7, Biolegend), Streptavidin (BV605, BD Biosciences). Data was acquired using the Aria Fusion (BD Biosciences) using BD FACSDIVA and analyzed using FlowJo (Beckon Dickinson).

### Quantitative Real-Time PCR

Approximately 3,000 cells were sorted directly into 400 µL of TRIzole (ThermoFisher #15596026), and RNA was isolated according to the manufacturer’s instructions. The RNA was reverse-transcribed using a SuperScript IV VILO Master Mix with ezDNase Enzyme kit (ThermoFisher #11756050). Quantitative Real-Time PCR was performed using PowerUp SYBR Green Master Mix (ThermoFisher #A25742) with a Quant Studio 4 (Applied Biosystems). qPCR primers are listed in Supplemental Table 1.

### Hydroxyproline Assay

Hydroxyproline was measured as previously described. The left lobe was homogenized using a Tissue Tearer (Biospec Products #985370-395), precipitated with trichloroacetic acid, baked in hydrochloric acid at 110 degrees Celsius overnight, reconstituted in water, then analyzed by colorimetric chloramine T assay as previously described (23).

### Precision Cut Human Lung Slices

The studies described in this paper were conducted according to the principles of the Declaration of Helsinki. Written informed consent was obtained from all subjects, and the study was approved by the University of California, San Francisco Institutional Review Board. Fibrotic lung tissues were obtained at the time of lung transplantation from patients with a diagnosis of Idiopathic Pulmonary Fibrosis or Silicosis. Basilar subpleural lung tissue was washed in PBS for 10 minutes x3. 18-gauge needles were used to instill 2% agarose (FisherScientific #BP1360-100) in PBS into the large airways, then left on ice for 30 minutes to solidify. The inflated lung was cut into approximately 2 cm^3^ cubes, mounted onto the stage of a Leica VT1200 vibratome. 500 µm thick sections were cut at 0.70 mm/s, then cut with scissors into approximately 1 cm height by 1 cm width slices and placed into a 24 well plate in PBS on ice. Lung slices were then transferred into new 24 well plates containing DMEM/F12+GlutaMAX (ThermoFisher #10565018) supplemented with Fungizone (ThermoFisher #15290026, 1:400), Primocin (InvivoGen #ant-pm-1, 100 µg/mL), Insulin-Transferrin-Selenium (Fisher Scientific #41-400-045, 1:100), EdU (ThermoFisher #A10044, 20 µM), and 1% Fetal Bovine Serum. Media was changed daily. Lung slices were cultured for 2 days at 37 degrees Celsius, 8% CO_2_, then washed in PBS and diced using scissors. Three lung slices were pooled into a single technical replicate. Three technical replicates (9 lung slices total) per donor were analyzed by flow cytometry then averaged into a single value for plotting.

### Histology

The right lung was inflated with 4% PFA in PBS under constant pressure of 25 cm H_2_O, tied off, then fixed in 4% PFA in PBS overnight at 4 degrees Celsius. Mouse lungs were then washed in PBS for 1 hour. The right upper lobe was used for cleared whole lung imaging. For confocal imaging, the right lung was transferred into 30% sucrose in PBS overnight, then embedded in Tissue-Plus Optimal Cutting Temperature compound (Fisher Scientific #23-730-571) for cryosectioning.

### Confocal Imaging of Brainbow2.1 Confetti Mouse Lungs

100 µm thick cryosections of mouse lung were washed in PBS then transferred into CUBIC-L (TCI #T3740) and incubated at 37 degrees on an orbital shaker (180 RPM) for 24 hours. Lung sections were stained with DAPI in PBS for 1 hour at room temperature, transferred into a 24-well glass bottom plate (Cellvis #P24-1.5H-N), then immersed in CUBIC R+M (TCI #T3741) for 2 hours at room temperature. 50 µm thick z-stack optical sections were obtained using an inverted Leica Stellaris Confocal Microscope. DAPI, GFP, YFP, and RFP channels were acquired for Scube2-CreER-Brainbow2.1mouse lung sections. Cthrc1-CreER labels a small percentage of lineage^+^ cells at baseline, so we only quantified cytoplasmic YFP^+^ and cytoplasmic RFP^+^ clones with clear fibroblast morphology for Cthrc1-CreER; Brainbow2.1 mice. 3D maximum projection images were generated in Imaris Viewer 10.2.0.

### Cleared Whole Lung and Light Sheet Imaging

After fixation in 4% PFA, the right upper lobe was transferred into CUBIC-L (TCI #T3740) and incubated at 37 degrees on an orbital shaker (180 RPM) for 24 hours x3, then washed in PBS as previously described (23). The cleared right upper lobe was RI-matched in CUBIC R+M (TCI #T3741) for 36 hours at room temperature, then transferred into RI 1.520 Mounting Solution for Cubic R+ (TCI #M3294). Images were acquired on the Nikon AZ100 Light Sheet microscope, using RFP channel to detect tdTomato and GFP channel to detect autofluorescence. 3D maximum projection images were generated using Imaris Viewer 10.2.0.

### Single-Cell RNA Sequencing Analysis for Publicly Available Mouse Lung Dataset

We used our previously published scRNA-seq dataset of bleomycin-treated Scube2-CreER; Ai14 mouse lung fibroblasts (GSE210341) (23). Using Seurat v5, the proliferating fibroblast cluster was subsetted and reclustered using NormalizeData, FindVariableFeatures, RunFastMNN, RunUMAP (1:20 dimensions), FindNeighbors (1:20 dimensions), and FindClusters (0.3 resolution). Differentially expressed genes for each of the proliferating fibroblast clusters were identified using the FindMarkers function (min percent 0.25, logfc.threshold 0.25). tdTomato^+^ cells were defined by natural-log-normalized tdTomato expression level greater than 3.5 to quantify the percent of tdTomato^+^ cells per cluster as previously described (23). Scaled average gene expression was used to calculate Spearman correlation coefficients for each cell cluster and plotted using pheatmap. Ggplot was used to generate scatterplots of differentially expressed genes comparing Proliferating fibrotic fibroblasts (day 7) versus reference alveolar fibroblasts (all days) and Non-proliferating fibrotic fibroblasts (day 7) versus reference alveolar fibroblasts (all days). Day 7 was selected because this time point had the highest frequency of proliferating fibroblasts.

### Single-Cell RNA Sequencing Analysis for Publicly Available Human Lung Dataset

We used our previously published (23) combined scRNA-seq Seurat object of pathologic and alveolar fibroblasts derived from normal and fibrotic human lungs by Tsukui (GSE132771) (23), Adams (GSE147066) (73), and Haberman (GSE135893) (63). We excluded cells from the Adams (GSE147066) dataset because Adams et al. regressed out cell cycle genes. Cell cycle scores were then calculated using the cell cycle scoring package in Seurat v5. We observed a lower frequency of high cell cycle scores in cells derived from the Tsukui (GSE132771) dataset, which may be due to longer storage time of fresh tissue prior to tissue dissociation. Thus, we focused exclusively on cells derived from the Habermann (GSE135893) dataset. To define proliferating versus non-proliferating cells, we used the Density-Based Spatial Clustering of Applications with Noise (DBSCAN) algorithm to identify hierarchical clusters based on cell cycle scores (eps 0.037, minPts 3). The decision tree model (rpart package) was then used to identify optimal cutoffs of cell cycle scores to delineate proliferating from non-proliferating cells (Supplemental Figure 4).

## Figures

Figure 2A was partially created in BioRender. https://BioRender.com/b92n121

## Statistics

Unpaired parametric t-tests were used for single comparisons between two groups. 2-way ANOVA with Tukey’s correction was used for multiple comparisons between 3 or more groups.

## Study Approval

Animal studies were approved by IACUC review boards. Protocol for harvesting human fibrotic lung tissue was approved by the UCSF Committee on Human Research, and written informed consent was received prior to participation.

## Data Availability

The scRNA-seq analysis was performed on publicly available data from accessions GSE132771 and GSE135893.

## Author contributions

C.M. and D.S. conceived the studies, interpreted the data, and wrote the manuscript. C.M performed all the experiments. T.T. contributed Scube2-CreER and Cthrc1-CreER mice and provided feedback on experimental design, methods, and manuscript revisions. I.K provided feedback on experimental design and data interpretation. X.R. and W.Q. assisted with harvesting mouse lungs for hydroxyproline experiments. P.W. and M.M. procured human lung samples.

## Acknowledgements

This work was supported by NHLBI 5R01HL142568-04. We thank UCSF core facilities: Laboratory for Cell Analysis for FACS and microscopes, Nikon Imaging Center for microscopes, and CoLabs Initiative for scRNAseq.

