## Supplemental Figures for "Clonal Expansion of Alveolar Fibroblast Progeny Drives Pulmonary Fibrosis"

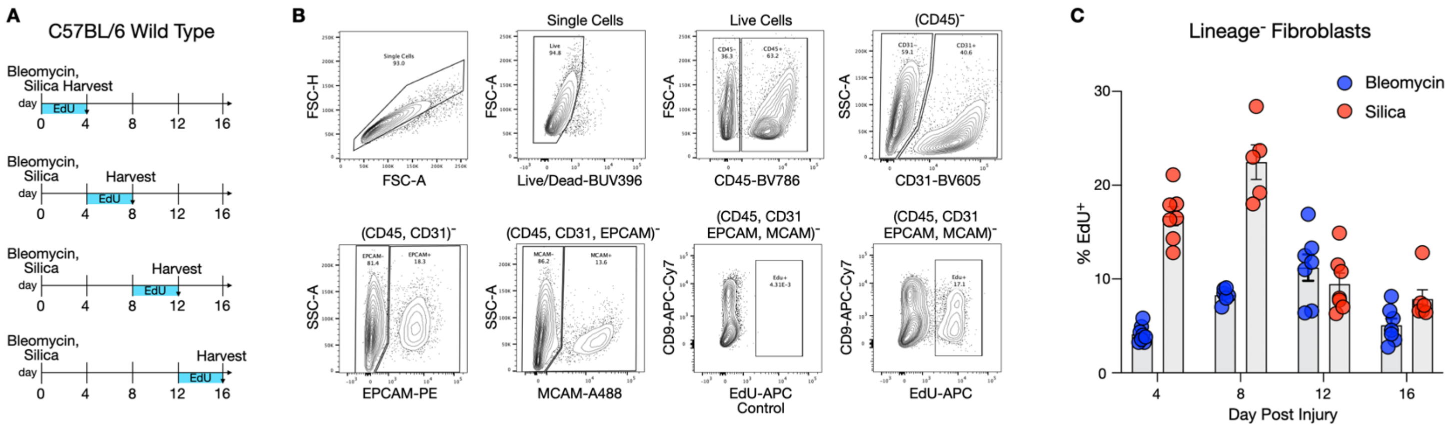


**Supplemental Figure 1: Proliferation Kinetics of Lineage- Fibroblasts in Fibrotic Lungs.** (**A**) C57BL/6 wild type mice were challenged with bleomycin or silica to induce pulmonary fibrosis then treated with daily injections of EdU for 4 days prior to harvest. Mice were harvested on days 4, 8, 12, and 16 post-injury and analyzed by flow cytometry. (**B**) Flow gating strategy for quantifying EdU uptake in lineage (CD45, CD31, EPCAM, MCAM)-negative lung fibroblasts. EdU-untreated mice were used as the EdU gating control. (**C**) Four-day average of % EdU uptake in lineage-fibroblasts after fibrotic lung injury with bleomycin or silica.

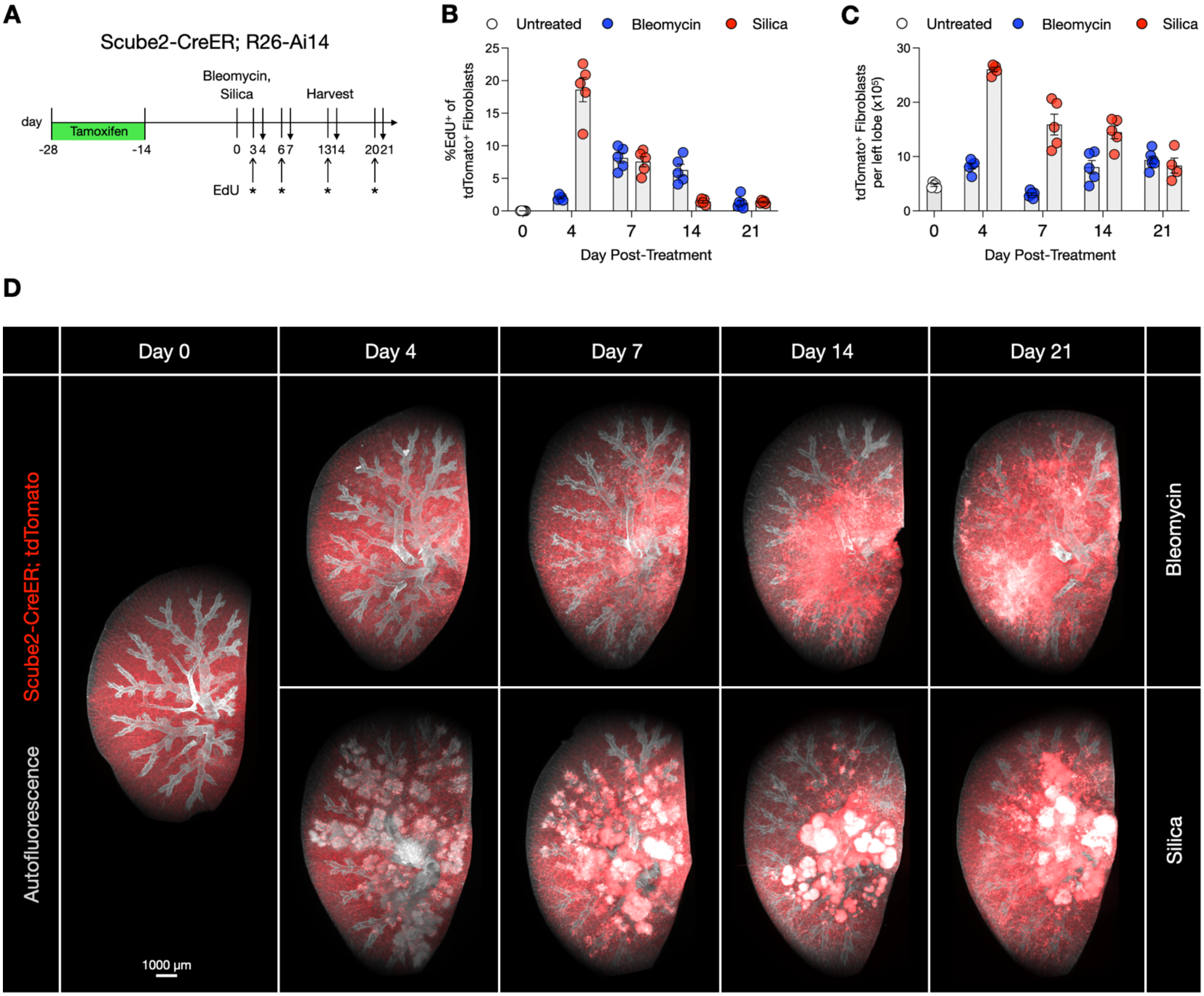


**Supplemental Figure 2: Proliferation Kinetics of Alveolar Fibroblast Descendants in Two In Vivo Models of Pulmonary Fibrosis.**
(**A**) Scube2-CreER/Rosa26-Ai14 mice were injected with tamoxifen to label alveolar fibroblasts with tdTomato, challenged with bleomycin or silica, then treated with EdU (*) 24 hours prior to harvest on days 4, 7, 14, and 21 post-injury. Total numbers of (CD45, CD31, MCAM, EPCAM)-negative tdTomato+ fibroblasts and their percentage of EdU uptake were quantified by flow cytometry. (**B**) Percentage of EdU uptake in tdTomato+ fibroblasts in bleomycin and silica-treated lungs, showing peak EdU uptake in silica-treated lungs at day 4 post-injury, and peak EdU uptake in bleomycin-treated lungs on day 7-post injury. (**C**) Quantification of absolute numbers of tdTomato+ fibroblasts post-injury. (**D**) 3D maximum projection images of cleared whole lungs showing the temporal emergence of tdTomato+ cell aggregates in bleomycin and silica-treated fibrotic lungs.

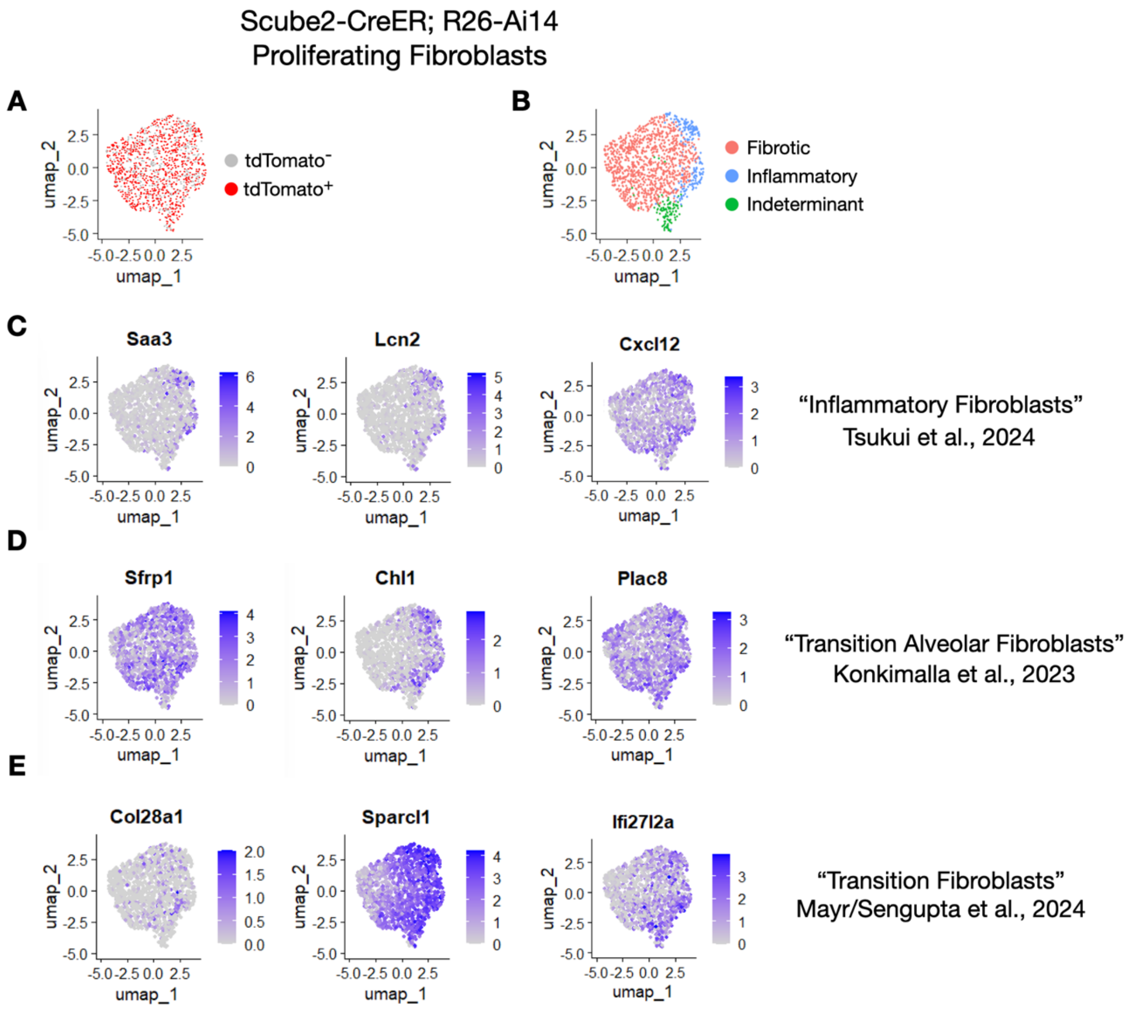


**Supplemental Figure 3: Fibroblast Subpopulation Comparison to Previous Studies.** (**A-B**) UMAP plot of proliferating fibroblasts from Figure 2, showing all subpopulations of proliferating fibroblasts were predominantly labeled by tdTomato. (C-D) Expression of genes related to previously described fibroblast subpopulations on UMAP feature plots of proliferating fibroblasts.

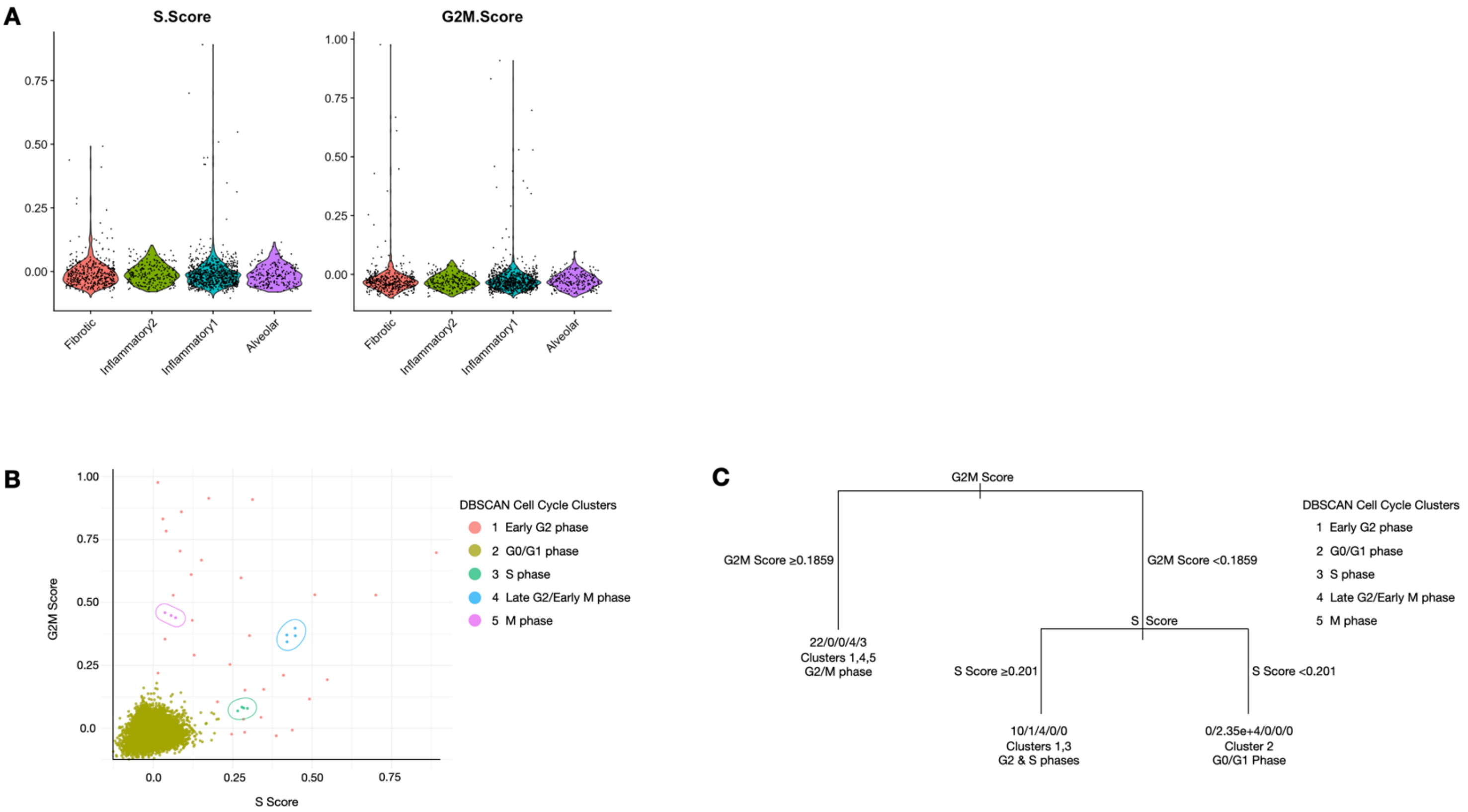


**Supplemental Figure 4: Identification of Proliferating Fibroblasts in Single Cell RNA Sequencing of Human Fibrotic Lungs.** Cell Cycle Scoring Seurat package was applied to fibroblast subsets in the re-analyzed Habermann et al. IPF Cell Atlas by Tsukui et al. 2024 (**A**) S phase and G2M phase cell cycle scores by fibroblast subtype. (**B**) Scatterplot of G2M and S phase cell cycle scores grouped into hierarchical clusters by the Density-Based Spatial Clustering of Applications with Noise (DBSCAN) algorithm. (**C**) Machine learning decision tree model identifies optimal cutoffs for S and G2/M phase scores to categorize cells as non-proliferative (DBSCAN Cluster 2) versus proliferative (DBSCAN Clusters 1,3,4,5). G2/M score ≥0.1859 and S score ≥0.201 were used to define cells in DBSCAN clusters 1,3,4,5 as "Proliferative", resulting in 99.99% of DBSCAN Cluster 2 (G0/G1) cells being defined as "Non-proliferative." Cell numbers per DBSCAN cluster are listed as #/#/#/#/# corresponding to DBSCAN clusters 1/2/3/4/5.

**Supplementary Table 1. qPCR Primers**

| | |
|---|---|
| Rps3 F | cggtgcagatttccaagaag |
| Rps3 R | ggacttcaactccagagtagcc |
| Scube2 F | CCTCTCTCAGAAGCAAACAGC |
| Scube2 R | GTCCTGACGGTGACGACAT |
| Pdgfra F | cggagcctgagctttgag |
| Pdgfra R | gccctgtgaggagacagc |
| Tcf21 F | CATTCACCCAGTCAACCTGA |
| Tcf21 R | CCACTTCCTTCAGGTCATTCTC |
| Npnt F | cagtgccaacctttctacgtc |
| Npnt R | tgtttgcactgtggttgaca |
| Cthrc1 F | aagcaaaaagcgctgatcc |
| Cthrc1 R | cctgctggtccttgtagacac |
| Col1a1 F | AGACATGTTCAGCTTTGTGGAC |
| Col1a1 R | GCAGCTGACTTCAGGGATG |
| Postn F | aagctgcggcaagacaag |
| Postn R | tcaaatctgcagcttcaagg |
| Tnc F | gggctatagaacaccgatgc |
| Tnc R | catttaagtttccaatttcaggttc |
